# Broad-spectrum mono- and combinational drug therapies for global viral pandemic preparedness

**DOI:** 10.1101/2022.01.15.476444

**Authors:** Aleksandr Ianevski, Rouan Yao, Ronja M. Simonsen, Vegard Myhre, Erlend Ravlo, Gerda D. Kaynova, Eva Zusinaite, Judith M. White, Stephen J. Polyak, Valentyn Oksenych, Marc P. Windisch, Qiuwei Pan, Eglė Lastauskienė, Astra Vitkauskienė, Algimantas Matukevičius, Tanel Tenson, Magnar Bjørås, Denis E. Kainov

**Affiliations:** Department of Clinical and Molecular Medicine (IKOM), Norwegian University of Science and Technology (NTNU), 7028 Trondheim, Norway; Vilnius Ozo gymnasium, Vilnius University, Vilnius 07171, Lithuania; Institute of Technology, University of Tartu, 50411 Tartu, Estonia; University of Virginia, Department of Cell Biology, Charlottesville, VA, USA; Virology Division, Department of Laboratory Medicine and Pathology, University of Washington, Seattle, WA, USA; Applied Molecular Virology Laboratory, Institut Pasteur Korea, 463-400 Gyeonggi-do, Korea; Department of Gastroenterology and Hepatology, Erasmus MC-University, Medical Center, Rotterdam, Netherlands; Department of Laboratory Medicine, Lithuanian University of Health Science, 44307 Kaunas, Lithuania; Life Sciences Center, Vilnius University, 10257 Vilnius, Lithuania; Institute for Molecular Medicine Finland, University of Helsinki, 00014, Helsinki, Finland

## Abstract

Broadly effective antiviral therapies must be developed to be ready for clinical trials, which should begin soon after the emergence of new life-threatening viruses. Here, we pave the way towards this goal by analyzing conserved druggable virus-host interactions, mechanisms of action and immunomodulatory properties of broad-spectrum antivirals (BSAs), routes of BSA delivery, and BSA interactions with other antivirals. Based on the analysis we developed scoring systems, which allowed us to predict novel BSAs and BSA-containing drug combinations (BCCs). Thus, we have developed a new strategy to broaden the spectrum of BSA indications and predict novel mono-and combinational therapies that can help better prepare for imminent future viral outbreaks.

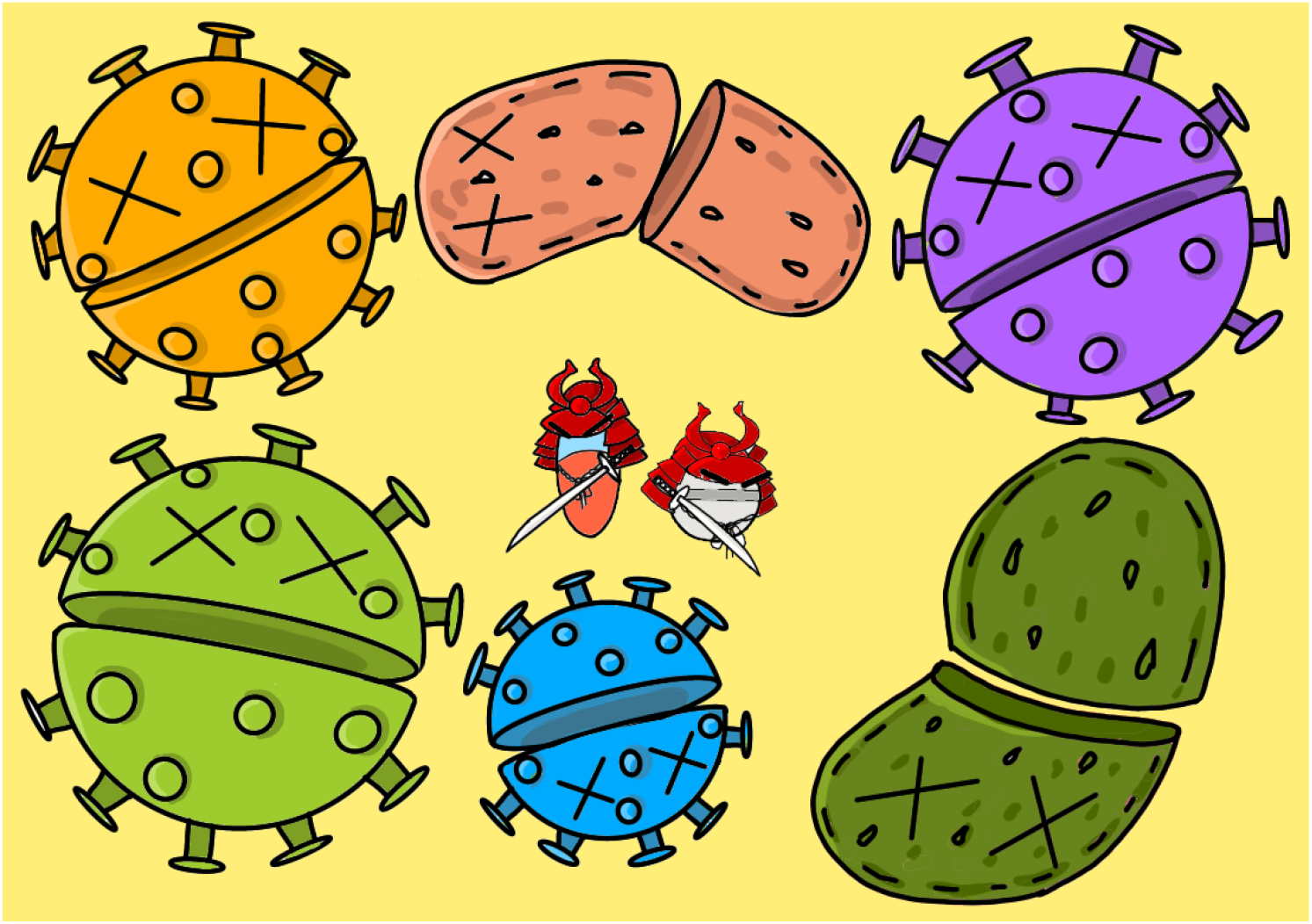

## Introduction

Despite advances in modern medicine, viral diseases still consistently pose a substantial economic and public health burden throughout the world. In fact, both the World Health Organization and the United Nations have highlighted the specific need for better management of viral diseases as priorities for future development [1]. This burden is likely due to viruses’ ability to regularly emerge and re-emerge into the human population from natural reservoirs such as wild and domesticated animals, leading to unpredictable outbreaks and wildly destructive health consequences [2]. However, despite this constant threat of viral outbreaks, the landscape of antiviral development is still underdeveloped, with over 200 human viral diseases that lack approved antiviral treatments.

Because the development of novel antivirals is long, laborious, and often unprofitable, the current strategy for management of viral outbreaks is heavily reliant on the development of vaccines over antiviral treatments [3]. However, while vaccines are an effective public health measure to stop community spread of a well-characterized virus, it is difficult to predictively develop vaccines against viral diseases that may emerge in the future. Therefore, antiviral development remains a crucial aspect of viral disease management to ensure timely and effective treatment of infected individuals and to reduce virus transmission.

Antiviral drugs are approved medicines that stop viruses from multiplying. Currently, there are 179 approved antiviral drugs, which are derived from 88 unique drug structures. Antiviral drugs currently represent 4.4% of 4051 approved medicines. However, 10 of 88 have been withdrawn due to side effects [4,5]. The most common side effects of many antiviral drugs are nausea, vomiting, allergic reactions, drowsiness, insomnia, heart problems and dependence [6]. Side effects can also be associated with the antiviral drugs that modulate the immune system, and may either enhance or suppress intrinsic immune functions of infected cells or alter the activity of immune cells within the host [7]. Antiviral drugs with immunostimulatory properties could lead to “cytokine storm”, which could be associated with an overwhelming systemic inflammation which leads to multiple organ dysfunction and potentially death [8]. By contrast, antivirals with immunosuppressive properties can be beneficial for the treatment of “cytokine storm” during influenza A virus (FLUAV) and severe acute respiratory syndrome coronavirus 2 (SARS-CoV-2) infections [9]. However, these drugs could prevent the development of adaptive immune responses allowing re-infections with the same or similar virus strains. Thus, antivirals without immunomodulatory properties are likely to be beneficial for treatment of viral infections [10,11].

Antiviral agents are molecules that have undergone pre-clinical development or clinical investigations against certain viruses but have not been approved for pharmaceutical use. Currently, there are thousands of antiviral agents in preclinical development and hundreds in clinical trials. It takes approximately 13-15 years and 2 billion USD to develop a new antiviral drug from an antiviral agent [12].

Antiviral drugs and agents can be further divided into those that target the virus and those that target the host. Virus-directed antivirals target viral proteins, viral nucleic acids, or lipid envelopes. An example of a virus-directed antiviral is oseltamivir, an influenza drug which inhibits viral neuraminidase. Host-directed antivirals target cellular factors that mediate virus replication. In contrast to virus-directed antivirals, host-directed agents modulate the activity of host factors and pathways. An example of host-directed antiviral is maraviroc, an HIV-1 drug which targets the cellular CCR5 receptor to prevent a critical step in HIV-1 entry.

Antiviral drugs and agents come in numerous molecular forms including small molecules, peptides, neutralizing antibodies, interferons (IFNs), Crispr-Cas systems, si/shRNAs, and other nucleic acid polymers (NAPs) [13-18]. Of these, neutralizing antibodies, peptides, NAPs, and Crispr/Cas are mainly used as virus-directed interventions; IFNs are used as host-directed drugs, while small molecules can be either virus-or host-directed drugs.

Broad-spectrum antivirals (BSAs) can inhibit replication of multiple viruses from different viral families [18]. One efficient method of BSA development is drug repurposing (also called drug repositioning), a strategy for identifying new uses for approved or investigational antiviral drugs that are outside the scope of the original medical indication [19]. BSAs are cost-effective because the overall development cost can be distributed across many viral indications. Critically, robust BSA development fosters future pandemic preparedness because BSA activity facilitates enhanced coverage of newly emerged viruses.

Ongoing viral replication and prolonged exposure to certain drugs can lead to the selection of drug resistant viruses through mutations in viral proteins. For example, mutations in HCV proteins confer resistance to NS3-4A, NS5A and NS5B inhibitors [20]. To mitigate the development of antiviral drug resistance, antivirals are combined [21]. Additive, multiplicative, and synergistic drug combinations are more effective than monotherapies, allowing for successful treatments at lower dosage and reduction of harmful side effects. Indeed, a combination of IFN-a and ribavirin was the “gold standard” for the treatment of chronic HCV infection for more than a decade [22]. Furthermore, ribavirin-and IFN-a-containing combinations have been used against other viruses [23,24] (NCT04412863), suggesting that BSA-containing combinations (BCCs) can be used to target a broad range of viruses.

Care needs to be taken when finding the correct BCCs. Drugs with unique mechanisms of action (MoA) are often paired together to minimize side effects and maximize efficacy. Drugs with the same MoA, such as nucleoside/nucleotide analogs, cannot be taken together [25] because they compete, rather than produce synergy or additive effect. Such combinations could also have higher toxicity than monotherapies. In addition, drug antagonism can reduce the effectiveness of treatment and lead to an increased risk of virologic failure (failure to meet a specific target of antiviral drug treatment). Ideally, one wants the smallest number of drugs in cocktail due to the potential for increased toxicity and additional side effects with each additional drug [26].

Here we have summarized the available scientific and clinical information regarding BSAs and BCCs, and reviewed the basic principles behind activities of monotherapies and synergism of combinational therapies. We also describe an approach to predict novel BSAs and BCCs for treatment of the most dangerous human viruses with pandemic potential. The approach described herein could facilitate the development of cost-effective and lifesaving countermeasures to fight new viral outbreaks.

### The landscape of BSA activities can be expanded

We extensively reviewed published BSAs using PubMed.gov, ClinicalTrials.org and DrugBank.ca. From this, we have identified 255 approved, investigational and experimental BSAs that target 104 human viruses from 24 families (Fig. 1; Supplementary data). Recently, we have tested several experimental, investigational, and approved BSAs against different viruses. We identified novel activities for saliphenylhalamide, gemcitabine, obatoclax, SNS-032, flavopiridole, nelfinavir, salinomycin, amodiaquine, obatoclax, emetine, homoharringtonine, atovaquone and ciclesonide, dalbavancin, vemurafenib, MK-2206, ezetimibe, azacitidine, cyclosporine, minocycline, ritonavir, oritavancin, cidofovir, dibucaine, azithromycin, gefitinib, minocycline, pirlindole ivermectin, brequinar, homoharringtonine, azacytidine, itraconazole, lopinavir, nitazoxanide, umifenovir, sertraline, amodiaquine and aripiprazole [27-41]. These results suggest that the landscape of BSA activities is vast and that it can and should be further interrogated and expanded.

**Fig. 1.**
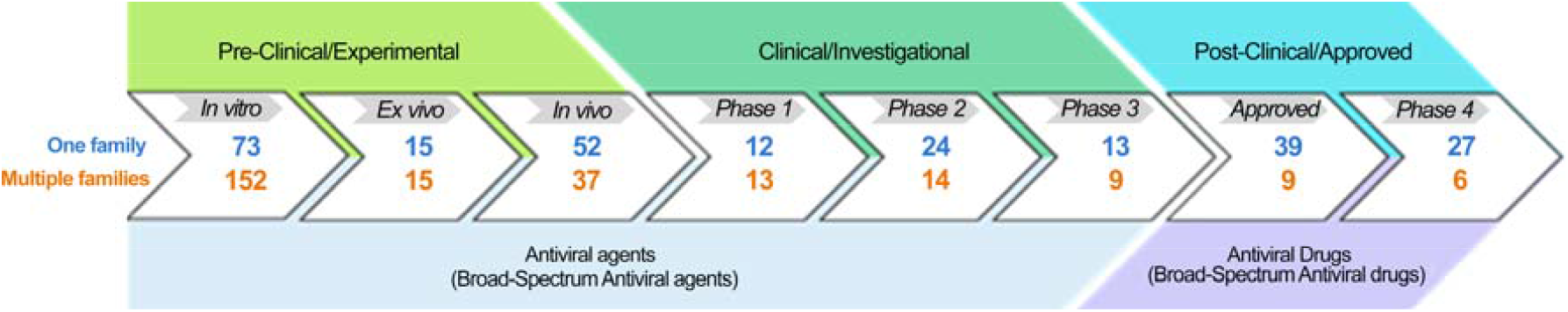
BSAs and stages of their development. BSAs that have reached the developmental stage in question for at least one viral indication are shown in blue, whereas BSAs that have reached the developmental stage in question for at least two viral indications are shown in orange.

To expand the activity spectrum of BSAs, we analyzed relationships between drug activity and virus phylogeny. For this, we first used the CLUSTALW2 algorithm [42] to build a phylogenetic tree based on available amino acid sequences of viral polymerases (pol) and reverse transcriptases (RT) (Fig. 2a). Then, we identified the number of BSAs found to have activity against each corresponding virus (Fig. 2b). Although the phylogenetically similar viruses will likely be responsive to the same drug, Fig. 2c indicates that most BSAs are only tested against a small subpopulation of related viruses. From this information, we can identify a wide range of previously untested BSA-virus interactions, which could demonstrate novel antiviral activities. For example, 54 BSAs have been proven effective for SARS-CoV-2, but not other coronaviruses such as SARS, MERS, 299E, NL63, OC43, or HKU1. From our analysis, we can infer higher likelihood of antiviral activity between those 54 BSAs and at least several other coronavirus species, strains or variants. Due to the high probability of coronavirus emergence, this inference could further be applied to coronaviruses that may arise in the future. However, it is important to note that our analysis here is limited to viruses which encode their own pols or RTs and for which full-length or near full-length sequences of these enzymes are available. For viruses that do not encode their own pols and RTs or are thus far poorly characterized, an analysis of virus taxonomy and BSA activity may be required to make similar inferences [43].

**Fig. 2.**
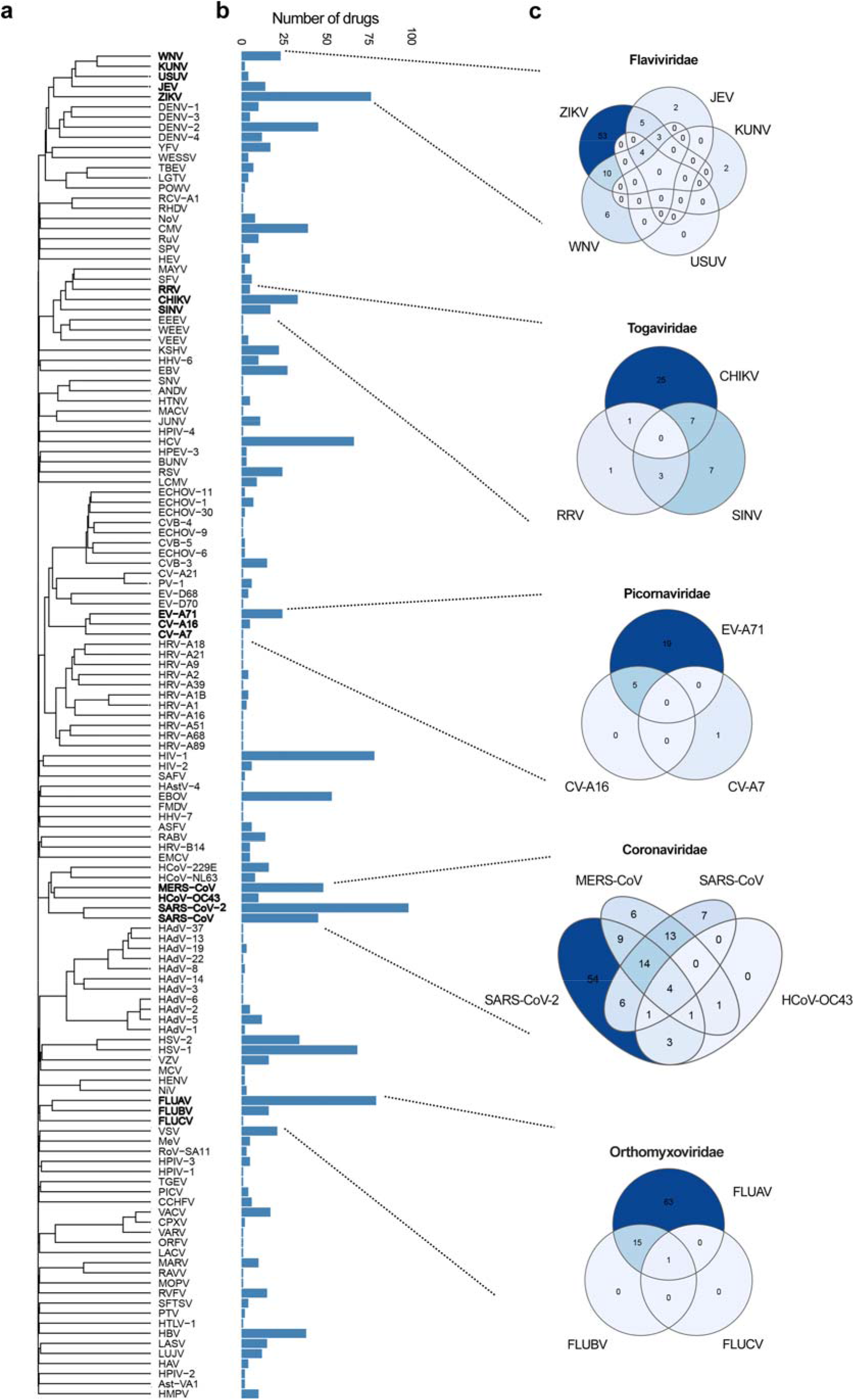
Drug activity-virus phylogeny relationship analysis. (**a**) Phylogenetic tree of viruses constructed based on the amino acid sequences of viral pols and RTs. (**b**) Bar chart showing the number of BSAs active against the viruses shown in panel (a). (**c**) Venn diagrams showing the number of BSAs targeting closely related viruses.

### Structure-activity relationship analysis identifies novel BSA candidates

To expand the list of potential BSAs, we performed drug structure-activity relationship (SAR) analysis of 11834 compounds from DrugBank 5.1.8 database [5], including 255 BSAs. Extended connectivity fingerprints of diameter 4 (ECFP4) were used to calculate the structural similarity of compounds [44]. We clustered compounds based on their structural similarities and extracted the compound sub-clusters that include two or more BSAs. Three such sub-clusters are shown in Fig 3. From this analysis, we can propose several new candidates for investigation as BSAs based on their structural similarities to existing BSAs. For example, the drugs domiphen, bephenium, pranlukast, afimoxifene, ospemifene, and fispemifene share structural similarity with tamoxifen and toremifene, known BSAs with activity against filoviruses and coronaviruses [45-48]. Based on structural similarity alone, we can identify these drugs as likely candidates for BSA activity. Similarly, pyronaridine, naphthoquine, meclinertant, and piperaquine are clustered together with BSAs quinacrine, amodiaquine, chloroquine, and hydroxychloroquine [49]; melarsomine, melarsoprol and FF-10101-01 are clustered together with BSAs etravirine, dapivirine and rilpivirine [50]; and CUDC-101, lapatinib, varlitinib, tucatinib, PD-168393, CP-724714, AZD-0424, tarloxotinib, canertinib, afatinib, dacomitinib, AV-412, falnidamol, enasidenib, LY-3200882, HM-43239, PD173955, PD-166326, abivertinib, olmutinib, poseltinib, spebrutinib, rociletinib, lazertinib, mobocertinib, osimertinib, alflutinib and TOP-1288 are clustered with BSAs erlotinib, saracatinib and gefitinib [51]. Thus, we demonstrate that this type of SAR analysis could identify critical BSA scaffolds and predict novel broad-spectrum antivirals.

**Fig. 3.**
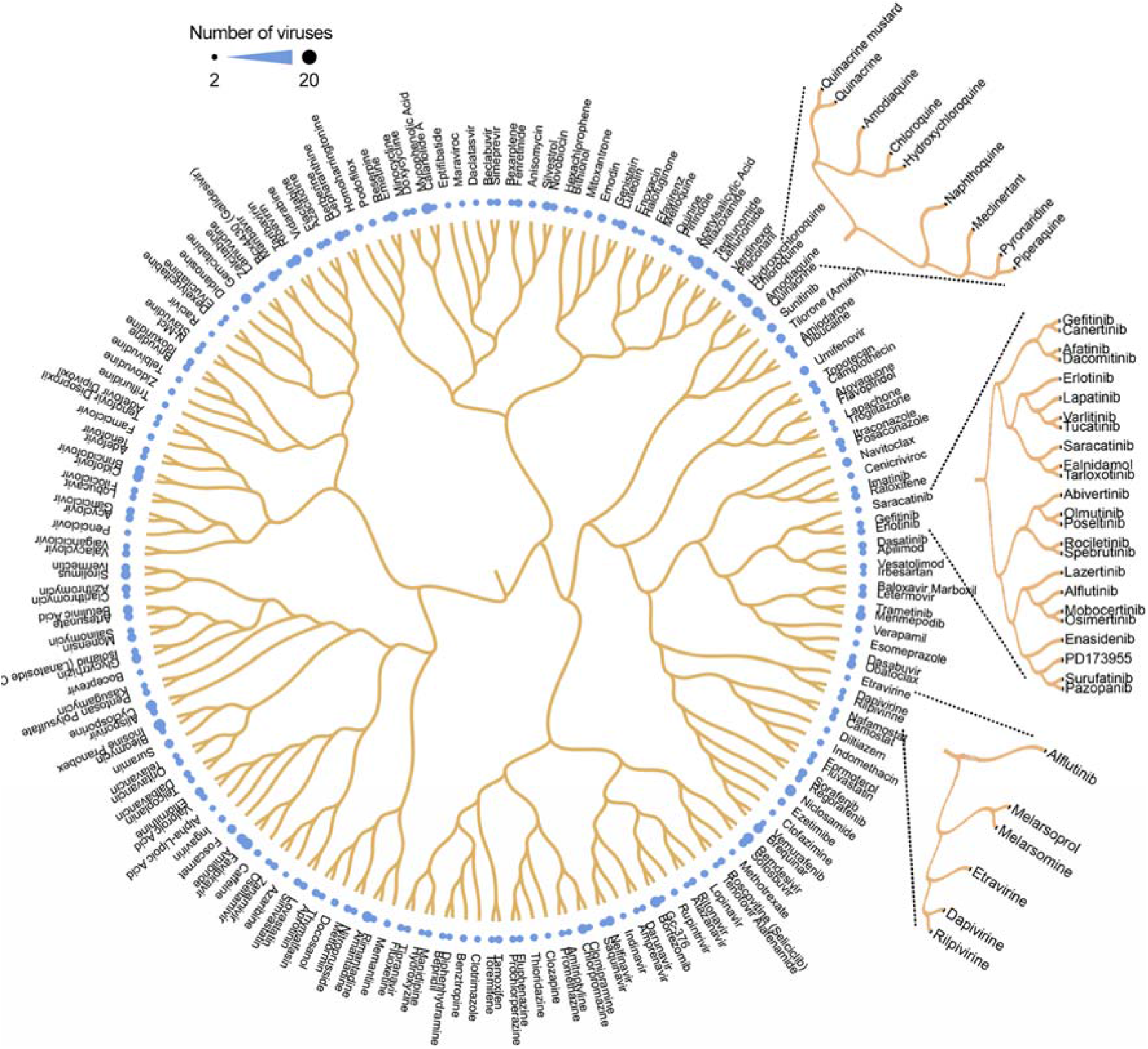
Structure-activity relationship analysis identifies compounds structurally similar to known BSAs. The circular dendrogram shows the SAR of BSAs from our database. We also used SAR analysis to identify BSA candidates from the list of 11834 compounds from DrugBank. Three compound sub-clusters that include two or more know BSAs are shown.

### Virus and host targets for BSAs

We next reviewed the known or suspected primary BSA targets (Supplementary data). We were able to identify primary targets for a fraction of BSAs. Most virus-directed BSAs work by inhibiting viral nucleic acid synthesis or protein processing (Fig. 4a). Among host-directed BSAs, mechanisms appeared to be more varied, and included inhibition of protein translation, trafficking, modification, or degradation, receptor-mediated signaling, lipid metabolism, etc. (Fig. 4b). However, in contrast to most host-directed BSAs that work through inhibition of host factors, several host-directed BSAs also work to activate innate immune responses against viruses. For example, IFNs are natural host-directed activators that bind their receptors to trigger cellular antiviral responses, which attenuate viral replication [52], and ABT-263 (navitoclax) targets the Bcl-xL protein to initiate apoptosis of infected cells without affecting non-infected cells [53]. Some BSAs, such as suramin, can simultaneously target host and viral factors [54,55]. Lastly, some BSAs are given in the form of prodrugs such as ganciclovir and gemcitabine which are activated by virus or host factors to achieve their antiviral effect [56].

**Fig. 4.**
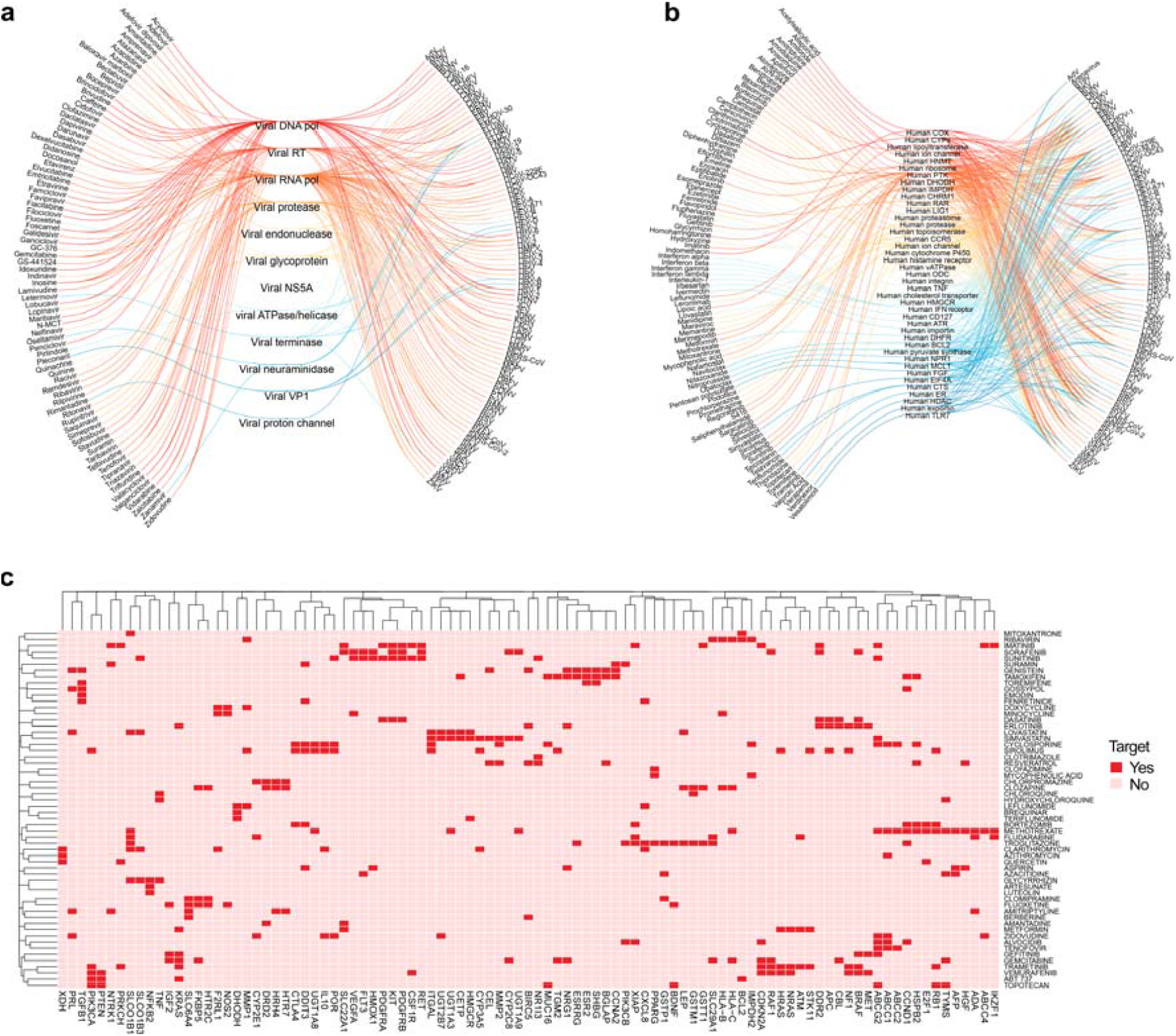
Virus and host targets for BSAs. (**a**) Eye diagram showing virus-directed BSAs linked to viruses through potential targets. (**b**) Eye diagram showing host-directed BSAs linked to viruses through potential targets. (**c**) Common targets of 58 BSAs, which possess immunomodulatory properties. Targets with interaction group scores <0.05 as well as unique targets were omitted. Clustering was performed to show highlight targets for BSAs.

By interrogating the Drug-Gene Interaction database [8], we found that several host-directed BSAs can target multiple cellular factors involved at several stages of viral replication. These extra drug targets are generally unexplored, and are most likely associated with side effects, although they may also affect viral replication. Perhaps, the most important of these targets are immunomodulatory. We compiled the BSAs with secondary immunomodulatory targets and showed those in Fig. 4c. From this, we found that many BSAs target similar clusters of immunomodulatory genes, indicating some structural and functional similarities between the targets, but no overarching immunomodulatory targets that may suggest a contribution to antiviral activity. Further analysis is needed to elucidate the exact role that these immunomodulatory targets play in host pharmacodynamics or their contribution to antiviral activity. Notably, the potential side effects of BSAs that are immunomodulatory can be mitigated by lower dosage of drugs in synergistic combinations.

### BSA-containing drug combinations for treatment of viral infections

Despite demonstrated efficacy at the early stages of drug development, many antiviral monotherapies are often found to be ineffective in clinical settings [57]. Because of this, antiviral cocktails have increasingly become the focus of drug developers. Antiviral combinations have several benefits over monotherapies. Namely, they can prevent the development of drug-resistant strains by completely halting viral replication, an advantage rarely achieved with monotherapies. Further, drugs administered together as cocktails may achieve an expanded antiviral activity, allowing for the treatment of multiple types of viral infection at once [58]. Because of this, BCCs are favorable candidates for front line therapy against poorly characterized emerging viruses, re-emerging drug-resistant viral variants, and viral co-infections.

Indeed, BCCs have become a standard treatment of rapidly evolving viruses, such as HIV and HCV (www.drugs.com/drug-class/antiviral-combinations.html). These include triple and quadruple drug combinations such as abacavir/dolutegravir/lamivudine (Triumeq), darunavir/cobicistat/emtricitabine/tenofovir (Symtuza), ledipasvir/sofosbuvir (Harvoni), sofosbuvir/velpatasvir (Epclusa), and lopinavir/ritonavir (Kaletra). Furthermore, many dual drug combinations are now in clinical trials against SARS-CoV-2, HCV, HBV, HSV-1, and other viral infections [59]. In addition, many BCCs have been tested in vitro and in animal models [27,34,60-63]. These and other studies further demonstrate the potential for antiviral combinations for the treatment of emerging and re-emerging viral infections.

To underscore the potential benefits and provide an organized summary of known dual antiviral drug combinations, we manually reviewed scientific literature and patent applications and constructed a BCC database (Supplementary data). The database comprises 538 drug cocktails. It includes 612 unique drugs and covers 68 viruses. We were able to identify primary targets for 415 drugs (Fig. 5). Of these, we found that 211 BCCs have components that both primarily target viral factors, 74 have components that both primarily target host factors, and 130 BCCs in which one drug primarily targets the virus while the other primarily targets the host. We were not able to identify specific targets for 160 BCCs due to one or more BSA in the BCC having an unknown mechanism of action. We suspect that the overrepresentation of virus-virus and virus-host targeting BCCs is due to the fact that drugs that were developed to specifically target virus factors may be more successful in achieving a direct antiviral effect while minimizing severe side effects. Thus, virus-virus and virus-host targeting BCCs are superior to host-host BCCs in many ways, including the leveraging of antiviral synergism, reduction of toxicity. However host-host targeting BCCs have lower risk of drug resistance, and an expanded spectrum of antiviral activity.

**Fig. 5.**
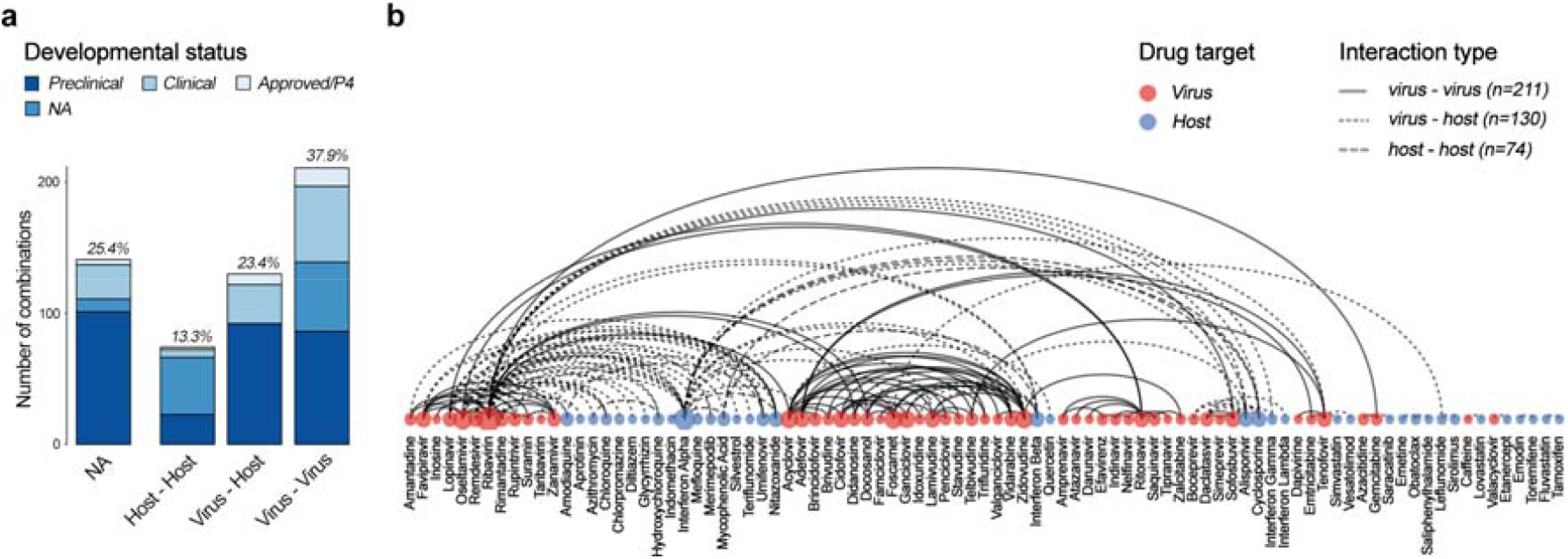
Drug-target interactions in BSA-containing combinations. (**a**) Developmental statuses and targets of BCCs. (**b**) Examples of BCCs targeting virus, host, or both factors. A random walks algorithm was used to group the drug combinations based on their targets [64].

### Association between infected organ systems and routes of administration of BSAs and BCCs

Viruses often preferentially infect hosts in one or more specific organ systems of the human body (Fig. 6a). In theory, BSAs and BCCs must be rapidly delivered to the infected organs using an amenable route of administration (RoA) to preserve the drug structure, maximize antiviral effect, and reduce drug toxicity or other adverse events. For example, if a virus infects and replicates in the respiratory system, medications administered by inhalation may be preferable. Likewise, if the virus infects the cardiovascular system, intravenous drug administration could be considered, etc. However, intravenous administration prevents widespread use of the BCCs because use is restricted to specialized care centers such as hospitals. In cases of advanced or systemic virus infections that affect multiple organ systems, antivirals intravenous administration may be preferable. However, most of the BSAs and BCCs reviewed here are delivered orally, most likely due to the preferential development of orally bioavailable drugs by pharmaceutical companies because of their increased marketability and potential for global distribution (Fig. 6b, c).

**Fig. 6.**
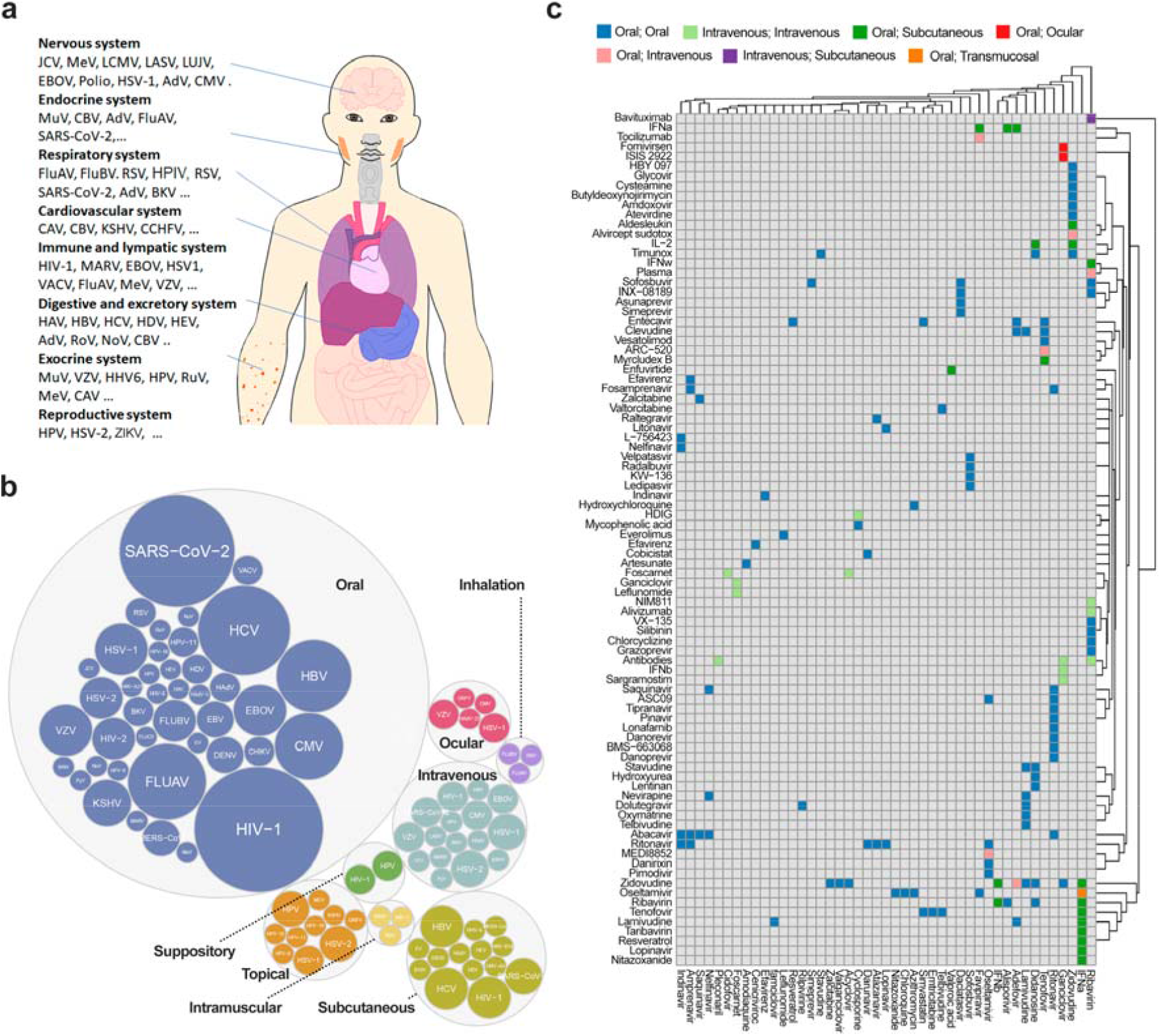
Routes of administration (RoA) of BSAs and BCCs. (**a**) Organ systems which are preferentially affected by different viruses. (**b**) RoA of BSAs. Sizes of the colored bubbles reflect the number of BSAs developed against a certain virus. (**c**) RoA of BCCs. Colored squares indicate the combined RoA of drugs in BCCs. Grey shading indicates that antiviral activity has either not been studied or reported for the drug combination in question.

### BSA and BCC scoring systems

To identify the most promising monotherapies we developed a six-component BSA scoring system:

1. SAR component (C_SAR_):
  - if the BSA is identical to a drug which has been developed or is currently under development for the virus of interest (voi), C_SAR_ = 1;
  - if the BSA is structurally similar to a drug which was developed or under development against the voi, C_SAR_ = 0.5;
  - if the BSA has a distinct structure, C_SAR_ = 0;
2. Drug developmental status component (C_DDS_; only applies to BSAs for which C_SAR_ = 1):
  - if the BSA is approved or is in phase 4 clinical trials against the Voi, C_DDS_ = 1;
  - If the BSA is in phase 1-3 clinical trials, C_DDS_ = 0.75;
  - if the BSA has been tested **in vivo**, C_DDS_ = 0.5;
  - if the BSA has been tested **in vitro**, C_DDS_ = 0.25;
  - - if the BSA has not been tested, C_DDS_ = 0;
3. Drug target relevance component (C_TR_):
  - if the confirmed primary target of the BSA in question is associated with Voi replication (the drug target is essential for Voi replication), C_TR_ = 1;
  - if not, C_TR_ = 0;
4. Drug immunomodulatory component (C_IC_):
  - if the BSA does not interfere with host immune response, C_IC_ = 1;
  - if the BSA is immunomodulatory, C_IC_ = 0;
5. Drug RoA component (C_RoA_):
  - if the RoA of the BSA is well-suited for the diseased system (for example, inhalation of drug for treatment of respiratory viruses), C_RoA_ = 1;
  - if not, C_RoA_ = 0;
6. Phylogeny component (C_Phyl_):
  - if the Voi is in the same genus as the virus for which the BSA has been developed, C_Phyl_ = 1;
  - if the Voi is in the same family, C_Phyl_ = 0.5;
  - if the Voi is in a closely-related family, C_Phyl_ = 0.25;
  - if the Voi is distantly-related, C_Phyl_ = 0.

To calculate the final BSA score, we sum the points across all six components using the following formula:

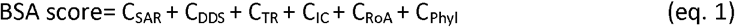

For example, the BSA score of favipiravir in relation to its activity against Ebola virus (EBOV) is 5.57. Favipiravir is an orally available nucleoside analog, which blocks viral RNA synthesis by inhibiting viral RdRP activity. Its immunomodulatory properties were not reported, and it is in phase 3 clinical trials against EBOV (NCT02329054). Therefore, the component values are as follows: C_SAR_=1, C_DDS_=0.75, C_DTR_=1, C_IC_=1, C_RoA_=1, and C_Phyl_=1 (Supplementary data).

Another example is merimepodib. Its BSA score in relation to its EBOV activity is 4.25. Merimepodib is an orally available inhibitor of host inosine monophosphate dehydrogenase (IMPDH), which controls the intracellular guanine nucleotide levels that are required for viral RNA synthesis. It possesses anti-EBOV activity in vitro and suppresses host immunity [23,65]. Therefore, the component values are as follows: C_SAR_=1, C_DDS_=0.25, C_DTR_=1, C_IC_=0, C_RoA_=1, and C_Phyl_=1 (Supplementary data).

To identify the most promising combinational therapies, we invented a four-coefficient BCC scoring system. It utilizes the following BCC coefficients:

1. Drug interaction coefficient (k_DI_):
  - if the MoAs for each component of the combination are different, k_DI_ = 1;
  - if the MoAs are the same (for example, if both components are nucleoside analogues), k_DI_ = 0.5;
2. Drug-target interaction coefficient (k_DTI_):
  - if both BSA components target viral factors (the combination for which minimum side effects are expected), k_DTI_ = 1.2;
  - if one BSA targets a viral factor and one BSA targets a host factor, k_DTI_= 1.1;
  - if both BSA components target host factors (the combination for which maximum side effects are expected, k_DTI_ = 1;
3. Drug-targeted stage of replication cycle coefficient (k_DRS_):
  - if both BSA components target the same stage of the virus life cycle (entry, viral replication, or exit), k_DRS_ = 1.2;
  - if the BSA components target different stages of the viral life cycle, k_DRS_ = 1;
4. Drug RoA coefficient (k_RoA_):
  - if both BSA components can be administrated by the same route and if the RoA can be used for targeted delivery to the diseased system, k_RoA_ = 1.2;
  - if both BSA components can be administrated by the same route, but the RoA cannot be used for targeted delivery to the diseased system, k_RoA_ = 1;
  - if the two BSA components cannot be administrated via the same route, k_RoA_ = 0.8; From these we calculate a BCC score using the following formula:

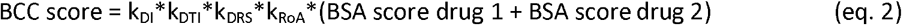

If the BCC score exceeds the sum of the individual BSA scores by 5, we consider this combination to be effective (Supplementary data). For example, for favipiravir-merimepodib targeting EBOV, the k_DI_ is 1.0 because the MoAs of the drugs are different; the k_DTI_ is 1.1, because the drugs target viral RdRP and host IMPDH; the k_DRS_ is 1.2, because both drugs reduce synthesis of viral RNA, and k_RoA_ 1.2, because both drugs can be taken orally, which allows delivery of the combination to multiple infected organs. Therefore, the BCC score of favipiravir-merimepodib is 15.8, whereas the combined BSA score of the combination is 10 (Table 1). Because the BCC score is greater than the combined BSA score by over 5 points, this combination would be considered to have high potential based on our scoring system. Indeed, by reviewing the literature, we found that this combination has been independently tested against EBOV in vitro and was shown to be effective [23].

**Table 1.**
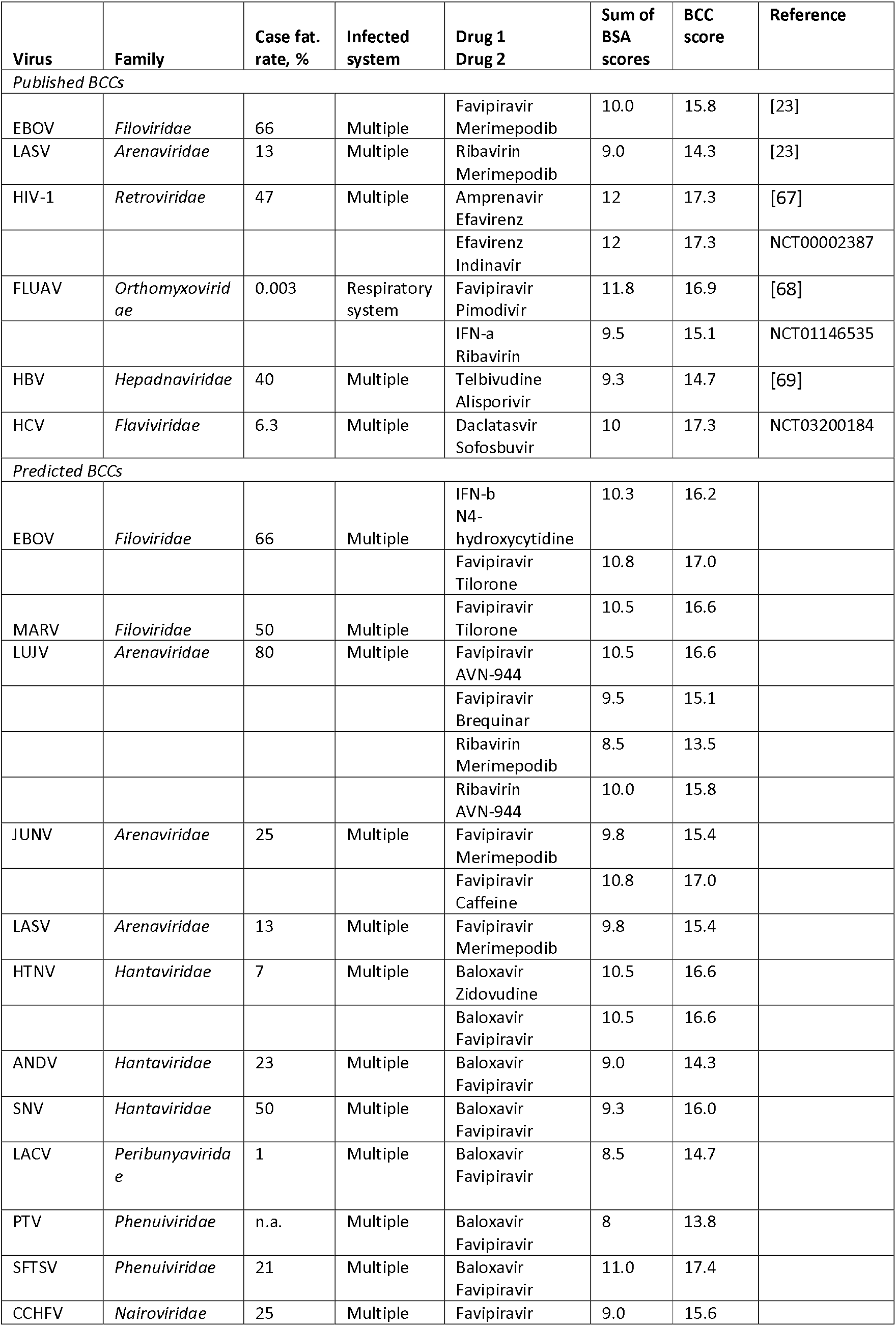
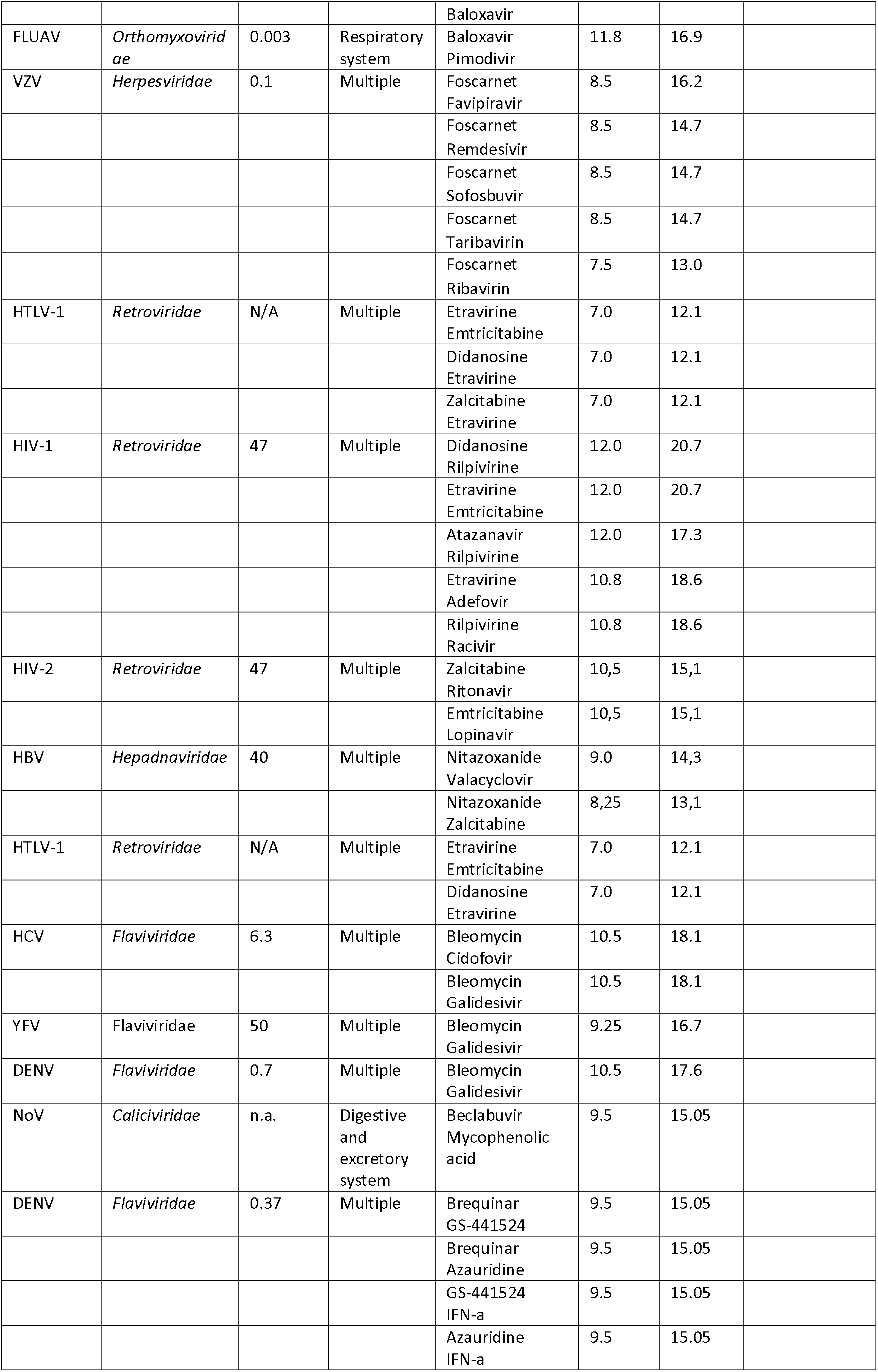
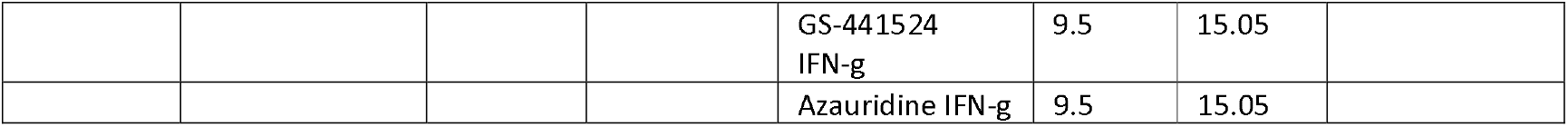
Examples of published and predicted BCCs, for which BCC scores exceed the sum of the individual BSA scores by 5.

In contrast, the BCC score for favipiravir-ribavirin against EBOV is 7.3. This is lower than the sum of individual BSA scores of favipiravir and ribavirin, which is 8.5, which predicts suboptimal performance for this combination (Table 1). Literature review shows that this prediction is consistent with efficacy studies in monkeys [66]. Thus, we demonstrated that the results from our scoring system are consistent with real-life experimental evidence.

Next, we used our scoring system for identification of novel potential BCCs (Supplementary data). As mentioned above, we focused on novel combinations for which BCC scores exceed the sum of the individual BSA scores by >5. In this way, we have identified several unexplored drug combinations that may be prioritized for development now in preparation for future resurgent outbreaks or the appearance of newly emerging viruses (Table 1). Thus, we identified novel BCCs for the potential treatment of life-threatening virus infections. Some of these combinations could be used as pan- and even cross-family BCCs. Analysis of scores of new and known BCCs could help prioritize the development of the best possible solutions.

### Conclusions and future perspectives

New life-threatening viruses emerge and pose a serious threat to public health. Thereby, broadly effective antiviral therapies must be developed to be ready for clinical trials, which should begin soon after a new virus start to spread from human to human [18]. To identify pan- and cross-virus family treatments, we established a scoring method, which is based on analysis of conserved druggable virus-host interactions, mechanisms of actions and immunomodulatory properties of BSAs, routes of their delivery, and BSA interactions with other antivirals. The method prioritizes the development of the most promising few of the thousands of potentially viable BCCs.

We intend to validate our proposed BCCs in cell cultures and animal models to confirm their effectiveness and ultimately prepare them for clinical trials. In particular, we will test selected drug cocktails in cell cultures against several life-threatening viruses and calculate synergy indexes to precisely distinguish synergistic from additive and antagonistic BCCs. Mathematical models that consider in vitro drug potency, pharmacokinetic and pharmacodynamic properties, and viral kinetics will further support prioritization of BCCs for in vivo studies [21]. We will than evaluate toxicity and efficacy, the immunological properties, and routes of administrations of BCCs using harmonized methods. In addition to our effort, we will invite other researchers to validate our proposed BCCs and develop our method further using machine-learning algorithms. This would allow us to prepare the most promising BCCs for clinical trials. If handled correctly, the development of the right BCCs can have a global impact by enhancing preparedness for future viral outbreaks, filling the void between virus identification and vaccine development with life-saving countermeasures and improving the protection of the general population against emerging viral threats.

## Ethics approval and consent to participate

Not applicable

## Consent for publication

The publisher has the author’s permission to publish research findings.

## Availability of data and material

Data is available within the article or its supplementary materials

## Authors’ contributions

All authors contributed to the methodology, software, validation, formal analysis, investigation, resources, data curation, writing, and review and editing of the manuscript. D.K. conceptualized, supervised, and administrated the study. All authors have read and agreed to the published version of the manuscript.

## Acknowledgments

We thank all the researchers and clinicians developing BSAs and BCCs.

